# Genetic Gains from Sixty Years of Spring Wheat Breeding in the Northern Plains of the US

**DOI:** 10.1101/2025.05.21.655386

**Authors:** Harsimardeep S. Gill, Sarah Blecha, Charlotte Brault, Karl Glover, Andrew Green, Jason Cook, Aaron Lorenz, Andrew Read, James A. Anderson

**Affiliations:** Department of Agronomy and Plant Genetics, University of Minnesota, St. Paul, MN 55108, USA; United States Department of Agriculture-Agriculture Research Service, Plant Science Research Unit, St. Paul, MN 55108, USA; Department of Agronomy, Horticulture and Plant Science, South Dakota State University, Brookings, SD 57007, USA; Department of Plant Science, North Dakota State University, Fargo, ND 58108, USA; Department of Plant Sciences and Plant Pathology, Montana State University, Bozeman, MT 59717, USA

## Abstract

Evaluating genetic gains over time is essential for assessing the success of breeding programs and refining strategies for ongoing improvement. Hard red spring (HRS) wheat is an important wheat class in the US and is primarily grown in the Northern Great Plains. Despite a long history of breeding efforts in this region, long-term quantification of genetic gains for key traits has remained limited. This study analyzes over sixty years of data from the USDA-coordinated Hard Red Spring Wheat Uniform Regional Nursery (HRSWURN) to evaluate genetic advancements in agronomic traits across multiple phases. A significant positive genetic gain of 0.61% per annum was observed for grain yield in HRS wheat released in the Northern US region, which is lower than the expected gains needed to meet future wheat demand. The change was 0.07% for test weight, −0.04% for days to heading, and −0.16% for plant height. Notably, sustained yield improvements have not affected grain protein levels since they were first measured in 1995, indicating that ongoing selection has effectively balanced grain yield and protein despite their negative correlation (r = −0.31). Assessment of genetic gains over 20-year phases suggested slowing rates of genetic gains for grain yield but did not indicate any plateaus. The realized genetic gains were generally higher for individual breeding programs when breeding for target environments, with the public breeding program in Minnesota observing gains of approximately 1% per annum. These findings highlight the significant impact of long-term breeding efforts and offer valuable insights for refining future breeding strategies.

## 1. Introduction

Wheat is a major staple crop with a rich history of breeding efforts focused on enhancing grain yield and quality. Wheat breeding has been instrumental in improving global food security through increased grain yields, improved resistance to pests, and better environmental resilience. The past century witnessed a Green Revolution resulting from improved high-yielding varieties combined with advanced agronomic practices that translated to substantial increases in wheat production, followed by continuous efforts to improve wheat yields. However, it has been suggested that gains in grain yield in recent decades have slowed or shown stagnation (Boehm et al., 2023; R. Fischer et al., 2014). In contrast, wheat production must increase by ∼ 50% to meet the expected demand by 2050 (Van Dijk et al., 2021). This requires continuous breeding efforts to ensure sustainable wheat production that meets projected demand.

Toward the goal of continuous improvement, it is important for a breeding program to measure the efficiency and track the genetic progress over long-term and short-term periods. Genetic trend, or realized rate of genetic gain, is considered one of the key indicators to measure the success of a breeding program (Ceccarelli, 2015). It is defined as an increase in performance expected or realized annually through selection for a given trait (Covarrubias-Pazaran, 2020; Rutkoski, 2019a, 2019b). Assessment of the rate of genetic gain not only provides an effective benchmark for the progress of the breeding program but also helps the program analyze its strengths and weaknesses and incorporate novel strategies adapted to new scenarios (Singh et al., 2007). For example, breeders employ different strategies and technologies to achieve specific breeding objectives. It could involve introducing more exotic germplasm to improve a specific trait or incorporating novel tools, such as genomic prediction or high-throughput phenotyping, into the breeding pipeline to accelerate the rate of cultivar development. A regular estimation of genetic gains allows breeders to determine the effectiveness of these technologies and make informed decisions to better optimize their breeding strategies (Covarrubias-Pazaran, 2020; Rutkoski, 2019a, 2019b; Seck et al., 2023; Xu et al., 2017).

Estimates of the genetic gain realized by a breeding program can simply be obtained by regressing the mean breeding value for a given trait on the year of origin (Eberhart, 1964; Rutkoski, 2019a). One of the most common methods is the era trial (Duvick, 2005), which involves pooling varieties developed by the breeding program over a time period and evaluating them within a single trial across multiple environments to obtain the mean breeding values. However, the ERA trials are prone to various logistical challenges and lack the ability for real-time monitoring and timely adjustments to breeding strategies (Menkir et al., 2022; Prasanna et al., 2022). Another useful approach is the use of historical datasets obtained from the evaluation of breeding lines over the years to estimate genetic gains (Mackay et al., 2011; Piepho et al., 2014). This approach enables periodic assessment and has been successfully employed in various crop breeding programs (Khanna et al., 2024; Menkir et al., 2022; Piepho et al., 2014; Seck et al., 2023). Nevertheless, this approach often suffers from a lack of connectivity between years or locations, which confounds the separation of genetic and non-genetic effects. To overcome this problem, it is recommended to use historical datasets connected through long-term checks to control the confounding of genetic and year effects, or make use of pedigree or genomic relationship matrices while obtaining the breeding values (Rutkoski, 2019a). Both of these approaches have been successfully utilized to estimate genetic gains across various crops, depending on the availability of resources or datasets (Asfaw et al., 2024; Ayenew et al., 2021; Khanna et al., 2024; Kumar et al., 2021; Mackay et al., 2011; Prasanna et al., 2022).

In wheat, long-term genetic trends have been extensively documented in different regions of the world to quantify genetic progress and inform future breeding strategies (Brancourt-Hulmel et al., 2003; Crespo-Herrera et al., 2018; Gerard et al., 2020; Gummadov et al., 2015; Sanchez-Garcia et al., 2013; Yadav et al., 2021). These reports include several studies from the US, but the majority represent winter wheat classes from different regions of the country (Battenfield et al., 2013; Boehm et al., 2023; Cox et al., 1988; Fufa et al., 2005; Graybosch & James Peterson, 2012; Graybosch & Peterson, 2010; Rife et al., 2019). Hard red spring (HRS) wheat is an important class of wheat grown in the US, accounting for approximately 25% of the country’s total wheat production and primarily grown in the Northern Great Plains ( USDA ERS, 2023). The HRS wheat breeding in the Northern Plains of the United States has a rich history, dating back to the late 19^th^ and early 20^th^ centuries, when breeding efforts were initiated to address the challenges posed by the region’s harsh climate and disease pressures (Paulsen & Shroyer, 2008). The era of early innovation was followed by the growth of public breeding programs and private seed companies, which expanded the scope of wheat improvement in the region. A significant milestone occurred in the 1960s with the introduction of semidwarf wheats. Since then, breeders have consistently developed varieties with enhanced agronomic traits, including improved yield potential, resistance to pests and diseases, and adaptability to the challenging conditions of the Northern Plains. Despite nearly a century of breeding efforts supported by public and private investments, no retrospective studies have been performed to monitor the rate of genetic progress in HRS wheat from the Northern US and Canadian prairies. A study to quantify yield gains, especially in the semidwarf era, would be beneficial for tracking the overall progress of HRS breeding in the region. In addition, different public breeding programs based on the land grant system have been developing cultivars for specific target populations of environments (TPE). The long-term and short-term genetic trends could be useful for individual programs to monitor their performance when breeding for their TPEs and develop new trait improvement strategies.

The Hard Red Spring Wheat Uniform Regional Nursery (HRSWURN), established about 90 years ago, has been instrumental in supporting spring wheat breeding efforts across the Northern Great Plains. The HRSWURN has undergone numerous name changes since its inception as a multi-state variety trial in 1929. Wheat breeders at land grant universities and private seed companies test advanced spring wheat lines each year in the HRSWURN for adaptability, yield, and disease resistance to determine if the entry should become a released cultivar for commercial production. The entries submitted by breeding programs are evaluated in replicated yield trials across multiple locations in the US and Canada (Figure 1). The historical HRSWURN dataset has been made publicly available, and some of the data have been successfully utilized in a study examining the impact of climate on wheat production (Zhang et al., 2022). However, no study has been performed to estimate genetic gains for HRS wheat using this historical dataset. More importantly, two long-term checks, ‘Marquis’ and ‘Chris’, have been part of the HRSWURN since 1968, providing an opportunity to use the historical data generated from this nursery to estimate genetic gains after the Green Revolution years. In the current study, we aim to assess the genetic gains for agronomic traits achieved over six decades of breeding efforts in the HRS wheat region using historical data from HRSWURN and to provide useful information to public HRS wheat breeding programs in North America.

**Figure 1.**
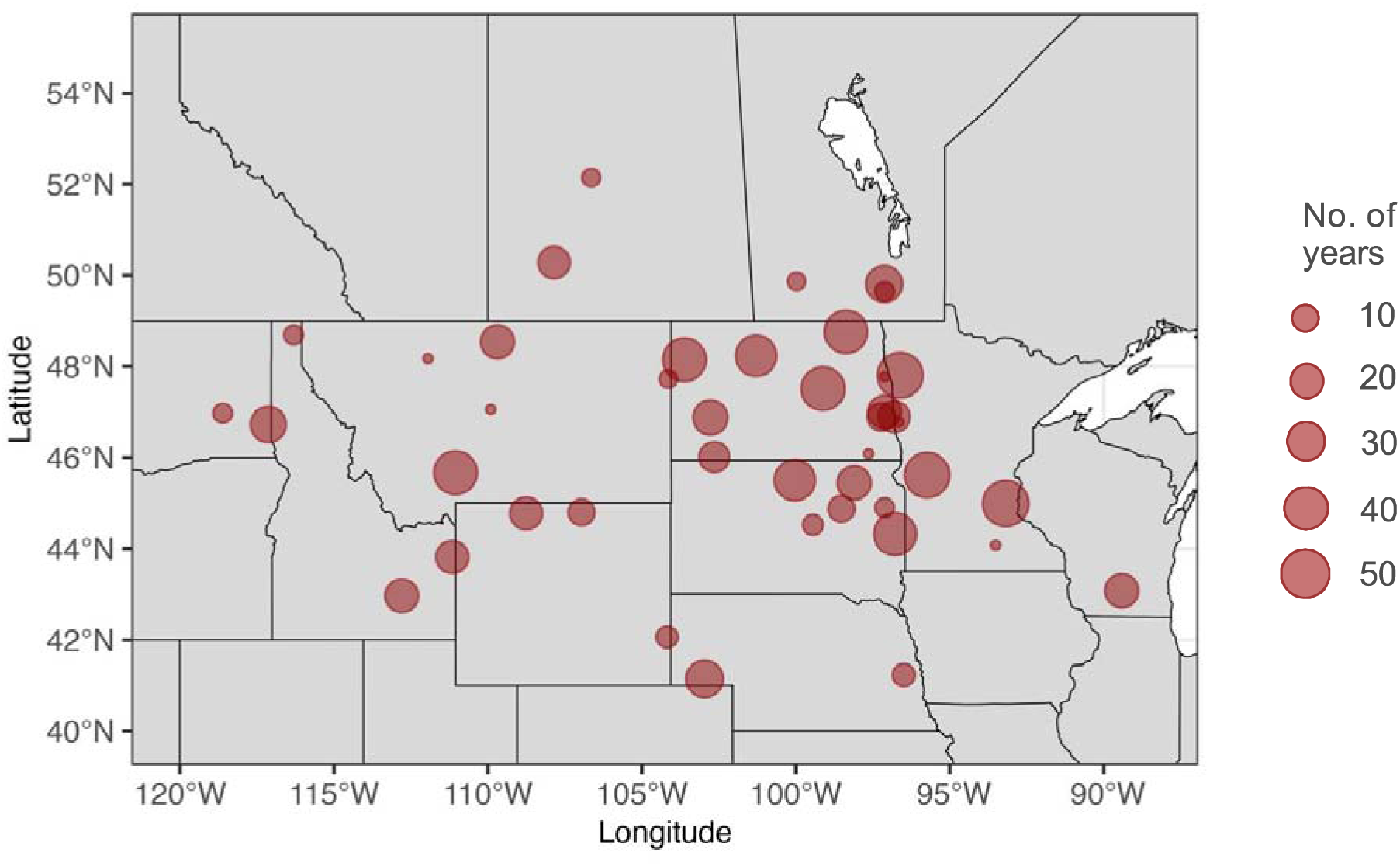
Geographic distribution and frequency of trial locations used in the Hard Red Spring Wheat Uniform Regional Nursery (HRSWURN) from 1968 to 2023. The map illustrates various testing locations across the Northern US and Canada used in HRSWURN trials over this period. Each point represents a unique location where trials were conducted. The circle size corresponds to the number of years a location was part of the nursery.

## 2. Materials and Methods

### 2.1 Description of the historical HRSWURN data

The HRSWURN is coordinated by scientists from the US Department of Agriculture-Agricultural Research Service (USDA-ARS), with various breeding programs contributing their advanced materials for multilocation trials. These replicated trials are conducted at diverse locations across several states, with the cooperators at the participating locations managing the trials, collecting and analyzing the data from their respective sites, and submitting it back to the coordinating scientist, who then compiles the multilocation data and USDA-ARS then publishes this data in the form of a comprehensive annual report. The annual reports for 1968-2023 are available in a digital format on GrainGenes, a publicly available data repository supported by the USDA-ARS (https://wheat.pw.usda.gov/GG3/). In addition to yield trials, the entries are evaluated for a variety of wheat diseases across different USDA disease-screening nurseries.

In this study, we utilized historical HRSWURN data spanning from 1968 to 2023, as it includes consistent long-term checks over this period. Although the starting year was 1968, a few varieties or breeding lines were either released or included in HRSWURN as early as 1960, which enabled us to estimate genetic gains over a six-decade period. Due to some logistical challenges, the HRSWURN was organized as a smaller ‘Tri-state nursery’ in 2011 and 2012, resulting in fewer trial locations during those years. The number of trial locations varied for each year, with 1987 having the highest number (22) and 2011 having the lowest (7) number of trial locations, resulting in a total of 1,026 unique environments (year-location combinations). The raw data were subjected to pre-processing and checked for any outliers or other remarks like hailstorms, drought, severe disease damage, etc. We excluded environments from the analysis based on remarks for growing challenges or high coefficient of variation (CV > 20%), leaving 957 environments for the final analysis in this study. Additionally, all the data were checked for any erroneous values, and the cleaned data were used for further analysis.

#### 2.1.1 Plant material and field design

The cleaned dataset from 1968 to 2023 comprised 1,016 unique genotypes contributed by various breeding programs. Most of these genotypes come from advanced stages of breeding programs, and some of these are later released as cultivars. For further analysis, we used the terms ‘all tested material’ and ‘released lines’ to differentiate these two categories. The unique genotypes also included two long-term checks (‘Marquis’ and ‘Chris’), as well as 11 short-term checks that were replaced periodically. The genotypes were primarily contributed by US land-grant universities/USDA breeding programs, private breeding programs, and Canadian public breeding programs (Supplementary Table S1). Among the public breeding programs, North Dakota (ND) contributed the highest number of unique genotypes (n = 273), followed by Minnesota (MN) (n = 213) and South Dakota (SD) (n = 157), and the frequency of test entries contributed by each program varied annually (Figure 2). Overall, the number of unique entries per year ranged from 21 in 1971 to 42 in 2008, with an average of about 32 entries per year. At each location, the trial was conducted using a randomized complete block design with three to four replications, and the plot size varied according to the testing environment.

**Figure 2.**
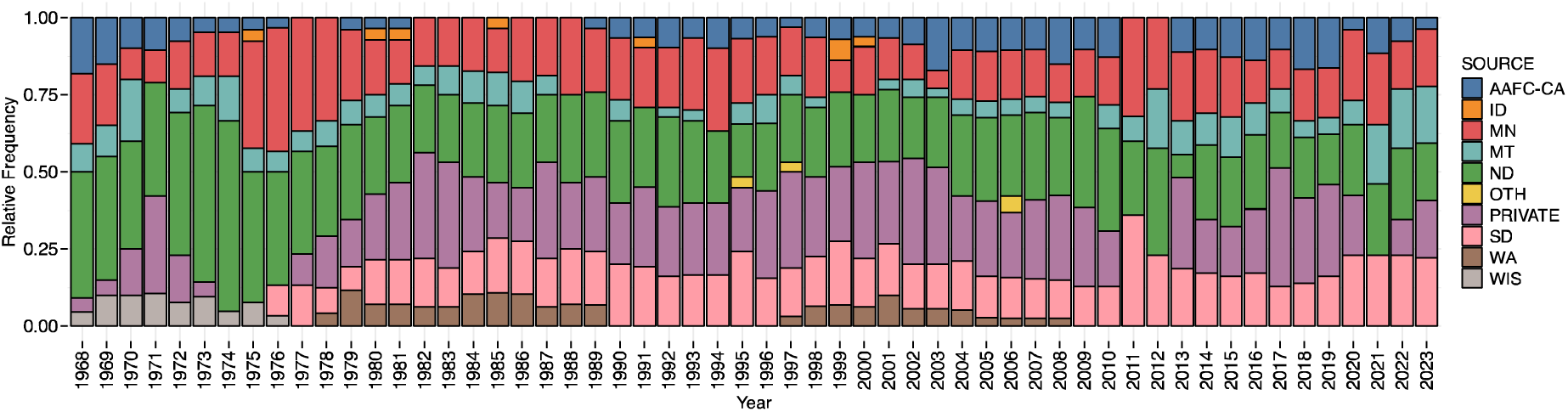
Sources of test entries in the Hard Red Spring Wheat Uniform Regional Nursery (HRSWURN) from 1968 to 2023. AAFC-CA, Canadian public breeding programs; ID, Idaho; MN, Minnesota; MT, Montana; ND, North Dakota; SD, South Dakota; WA, Washington; WIS, Wisconsin; OTH, Other public breeding programs with few entries. PRIVATE refers to the lines contributed by different seed companies from the private sector.

#### 2.1.3 Phenotyping and data collection

The phenotypic evaluation of the HRSWURN entries varied by the trial location. The major agronomic traits included grain yield (YLD), test weight (TWT), plant height (HT), days to heading (HD), lodging (LODG), thousand kernel weight (TKW), and grain protein content (PROT). Protein content was not available before the 1995 nursery. Many locations did not report all the agronomic traits (Figure 3; Supplementary Figure S1). In addition to the agronomic traits, several locations reported scores for wheat rusts (caused by *Puccinia spp*.) and Fusarium head blight (caused by *Fusarium graminearum*). For the current study, we focused on YLD, TWT, HT, HD, and PROT. As the data were reported in different units across years and locations, we standardized YLD to kilograms per hectare (kg ha^-1^), TWT to kilograms per hectoliter (kg hL^-1^), and HT to centimeters (cm). The HD was recorded as the number of days to heading as Julian days, and PROT was recorded as a percentage. After pre-processing, grain yield (YLD) was the trait available from the highest number of environments, followed by test weight (Table 1). The total number of observations available for YLD, TWT, HT, HD, and PROT was 29,422, 28,798, 27,983, 25,381, and 8,760, respectively. A detailed description of the number of genotypes evaluated for a respective trait across location-years is provided in Table 1.

**Figure 3.**
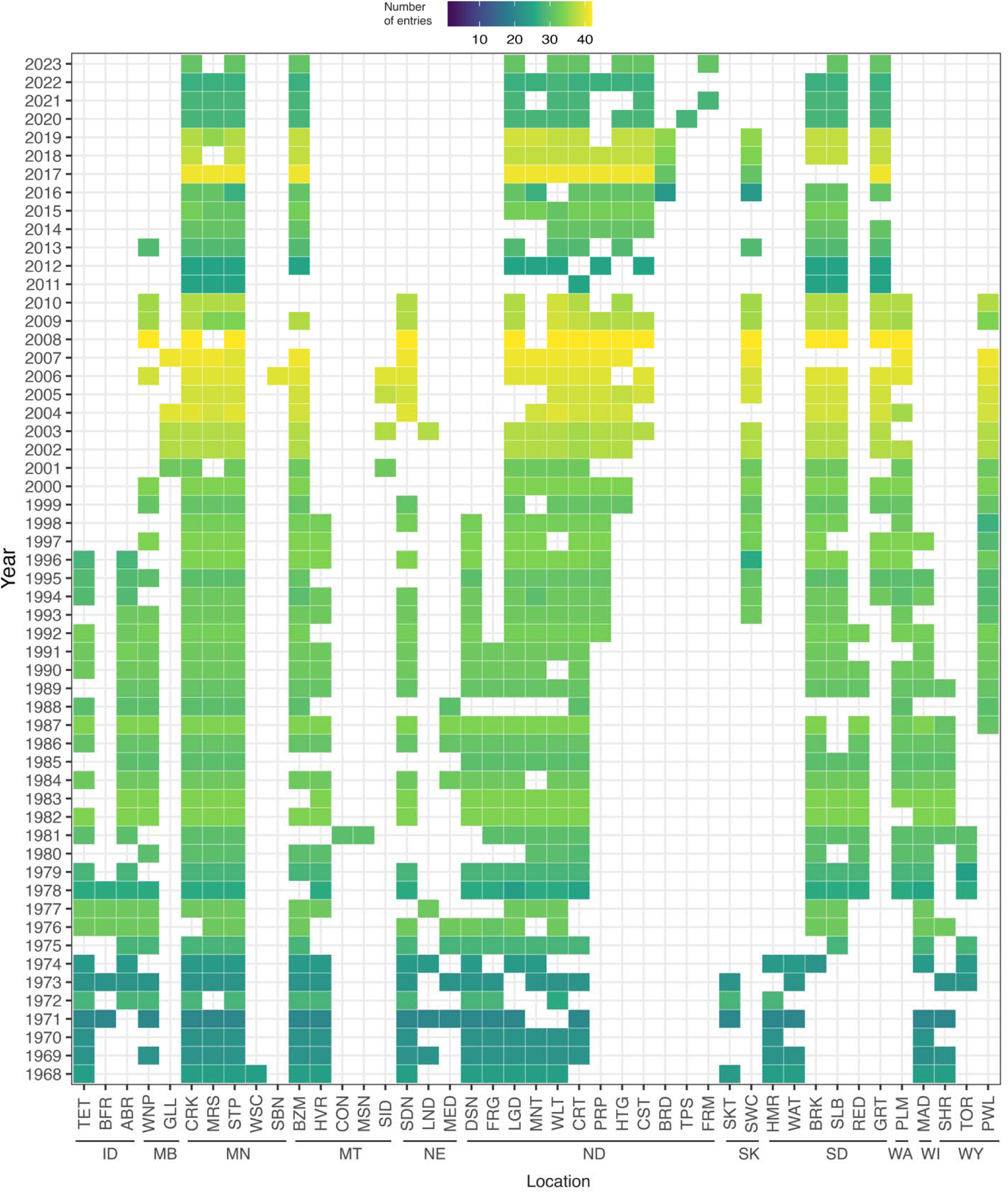
Locations based on the cleaned dataset where grain yield (YLD) was collected from 1968 to 2023. For a particular year and location, white indicates that data for YLD was not reported or the given location was not part of the nursery in the respective year. The scale provided in the legend elucidates the number of genotypes tested. The locations are nested under the State/Province with ID, Idaho; MB, Manitoba; MN, Minnesota; MT, Montana; NE, Nebraska; ND, North Dakota; SK, Saskatchewan; SD, South Dakota; WA, Washington; WI, Wisconsin; and WY, Wyoming. Detailed information and abbreviations for locations are provided in Supplementary Table S2.

**Table 1.** The number of genotypes evaluated in a given year and the number of locations where a respective trait was measured in a given year. Within each cell of the table, the number of genotypes is listed first, followed by the number of locations. The “–” symbol indicates that no data were collected.

| YEAR | YLD | TWT | HD | HT | PROT |
| --- | --- | --- | --- | --- | --- |
| 1968 | 24;18 | 24;17 | 24;14 | 24;16 | - |
| 1969 | 22;19 | 22;18 | 22;15 | 22;16 | - |
| 1970 | 22;15 | 22;15 | 22;13 | 22;15 | - |
| 1971 | 21;20 | 21;19 | 20;14 | 20;15 | - |
| 1972 | 28;13 | 28;13 | 28;11 | 28;11 | - |
| 1973 | 23;20 | 23;20 | 22;16 | 22;17 | - |
| 1974 | 23;17 | 23;17 | 23;15 | 23;13 | - |
| 1975 | 28;17 | 28;16 | 28;14 | 28;16 | - |
| 1976 | 32;17 | 32;17 | 32;9 | 32;14 | - |
| 1977 | 32;16 | 32;17 | 32;13 | 32;15 | - |
| 1978 | 26;21 | 26;21 | 26;16 | 26;21 | - |
| 1979 | 28;20 | 28;20 | 28;17 | 28;20 | - |
| 1980 | 30;14 | 30;14 | 30;13 | 30;14 | - |
| 1981 | 30;19 | 30;18 | 30;17 | 30;18 | - |
| 1982 | 34;20 | 34;20 | 34;17 | 34;20 | - |
| 1983 | 34;19 | 34;18 | 34;17 | 34;17 | - |
| 1984 | 31;21 | 31;20 | 31;19 | 31;20 | - |
| 1985 | 30;17 | 30;17 | 30;16 | 30;17 | - |
| 1986 | 31;21 | 31;21 | 31;18 | 31;19 | - |
| 1987 | 34;22 | 34;21 | 34;21 | 34;21 | - |
| 1988 | 30;11 | 30;10 | 30;8 | 30;10 | - |
| 1989 | 31;21 | 31;20 | 31;19 | 31;20 | - |
| 1990 | 32;20 | 32;19 | 32;18 | 32;20 | - |
| 1991 | 33;19 | 33;18 | 32;17 | 33;19 | - |
| 1992 | 33;20 | 33;19 | 33;18 | 33;21 | - |
| 1993 | 32;19 | 32;18 | 32;18 | 32;19 | - |
| 1994 | 32;21 | 32;21 | 32;19 | 32;20 | - |
| 1995 | 31;20 | 31;18 | 31;17 | 31;19 | 30;5 |
| 1996 | 34;20 | 34;20 | 34;19 | 34;20 | 34;4 |
| 1997 | 34;17 | 34;16 | 34;14 | 34;17 | 34;4 |
| 1998 | 34;18 | 34;17 | 34;16 | 34;18 | 33;4 |
| 1999 | 31;17 | 31;16 | 31;15 | 31;16 | 31;5 |
| 2000 | 34;17 | 34;16 | 34;15 | 34;17 | 34;6 |
| 2001 | 32;14 | 32;14 | 32;12 | 32;14 | 32;6 |
| 2002 | 37;17 | 37;17 | 37;14 | 37;17 | 37;9 |
| 2003 | 37;19 | 37;19 | 37;16 | 37;19 | 37;10 |
| 2004 | 40;16 | 40;16 | 40;15 | 40;16 | 40;10 |
| 2005 | 39;16 | 39;16 | 39;14 | 39;16 | 39;11 |
| 2006 | 40;20 | 40;20 | 40;18 | 40;20 | 40;11 |
| 2007 | 41;15 | 41;15 | 41;13 | 41;15 | 41;10 |
| 2008 | 42;15 | 42;15 | 42;13 | 42;15 | 42;10 |
| 2009 | 41;18 | 41;18 | 40;16 | 41;18 | 41;13 |
| 2010 | 41;15 | 41;15 | 39;13 | 41;14 | 41;11 |
| 2011 | 25;7 | 25;7 | 25;6 | 25;6 | 25;6 |
| 2012 | 26;12 | 26;12 | 26;11 | 26;12 | 25;7 |
| 2013 | 29;13 | 29;11 | 29;11 | 29;10 | 29;9 |
| 2014 | 31;12 | 31;12 | 31;12 | 31;12 | 31;10 |
| 2015 | 33;13 | 33;13 | 33;11 | 33;13 | 33;7 |
| 2016 | 31;15 | 31;15 | 31;13 | 31;15 | 31;10 |
| 2017 | 41;14 | 41;14 | 41;7 | 41;14 | 41;11 |
| 2018 | 38;15 | 38;16 | 38;13 | 38;16 | 38;13 |
| 2019 | 39;15 | 39;15 | 39;13 | 39;13 | 39;11 |
| 2020 | 28;14 | 28;14 | 28;14 | 28;14 | 28;11 |
| 2021 | 28;12 | 28;12 | 28;10 | 28;12 | 28;11 |
| 2022 | 28;14 | 28;14 | 28;13 | 28;14 | 28;12 |
| 2023 | 31;11 | 31;11 | 31;10 | - | 31;10 |

### 2.2 Statistical analysis

The HRSWURN trials are conducted at multiple locations across states and are hosted by individual breeding programs. The hosting programs manage the replicated trial, collect the data, and perform appropriate statistical analysis for individual locations to correct for experimental error. The adjusted entry means for each location, along with the CV for the trial (not for all locations), are reported to the USDA-ARS nursery coordinator and are compiled in the form of an annual report. In this study, we used the adjusted entry means for different traits from each location. A two-step method was followed to analyze the historical data over the years and estimate the realized genetic gains.

To estimate the mean performance of the genotypes across environments, a linear mixed model was used to obtain the BLUE (best linear unbiased estimate) for each genotype effect (Mackay et al., 2011; Raymond et al., 2023). The following model was fit to the data:

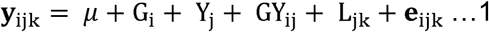

where y_ijk_ is adjusted entry mean of the i^th^ genotype in year j at site k obtained from the first stage of analysis, *μ* is the overall mean, G_i_ is the effect of the i^th^ genotype, Y_j_ is the effect of the j^th^ year, GY_ij_ is the genotype-by-year interaction effect, L_jk_ is the effect of location k nested within year j, and e_ijk_ is the residual, due to the combined effects of within-trial error and genotype × location within-year interaction. The genotype and year effects were both considered as fixed effects. According to Mackay et al. (2011), considering the genotype and year as random effects could produce bias. Hence, the above model has been widely used to estimate genetic gains in various crop species using historical datasets (Asfaw et al., 2024; Covarrubias-Pazaran, 2020; Delgado et al., 2024; Seck et al., 2023). Except for genotype and year, all other terms in Equation 1 were included as random effects. The broad-sense heritability of the estimated genotype effects using this highly unbalanced data was calculated by fitting the genotype term and its interaction with location as random effects in the above equation and using the formula described in Cullis et al. (2006):

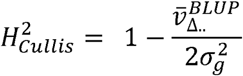

where 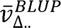 is the mean variance of the difference between two BLUPs for the genotype, which is the genotypic variance.

In the second step, realized genetic gains were estimated using a simple regression equation, where the trait BLUEs were regressed against the year of origin to obtain the rates of genetic gain per year, which were given by the slope of the linear regression line and indicated the annual change in the trait value. The intercept denoted the hypothetical value of the trait at the onset of the breeding program (year zero in the regression equation) (Delgado et al., 2024). For consistency, the year of origin was considered the first year any genotype was included in the HRSWURN. Though we used the HRSWURN data from 1968, there were a few genotypes in the initial years (1968-70) that were part of the HRSWURN from the 1960s. For these genotypes, we used the exact year they were included in the nursery, or the year they were released as a cultivar as the year of origin. In addition to this, the two checks (‘Marquis’ and ‘Chris’) were removed from the analysis while fitting the regression equation to avoid any bias in the genetic gains estimates, as suggested by Raymond et al (2023).

The percentage change in realized genetic gain was calculated for each trait, as suggested by Covarrubias-Pazaran (2020), from the ratio of the regression slope to the trait value at the first crossing year (year 1). This ratio showed the relative change/increase in trait values per year as follows:

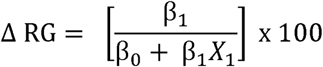

Where Δ RG is the percentage change in realized genetic gain calculated as the ratio of the slope of the regression model (β_1_) to the y-intercept of the regression (β_0_) plus the slope multiplied by the first year of origin.

To better understand the trends over time, we divided the dataset into three phases and estimated the genetic gains for grain yield in each of these phases using only released cultivars. The entire dataset was divided into three twenty-year phases to get an accurate estimate of genetic gains, as shorter time windows might be affected by various factors, including environmental pressures. The first phase, the pre-1984 period, marked the introduction and adoption of semidwarf wheats, as well as the implementation of improved management practices. The second phase, from 1984 to 2003, included further improvement in agronomics, the devastating Fusarium head blight epidemics in the spring wheat region, and the introduction of exotic germplasm for disease resistance. The third phase, from 2004 to 2023, represents the application of molecular marker technologies and integration of genomics and other modern tools into the breeding programs.

### 2.3 Genetic gains when breeding for target environments

In addition to estimating the genetic gains in HRS wheat from the Northern Region as a whole, we assessed the trends for grain yield improvement from individual public breeding programs based in Minnesota, North Dakota, and South Dakota. For this, the trial locations within the target population environment of individual wheat breeding programs were selected (Supplementary Table S2), and genetic gains were estimated using the method described earlier. For each of these programs, we used the breeding lines and released cultivars from the respective program only to estimate realized genetic gains.

### 2.4 Software and code

All statistical analyses were performed in the R environment using the base or other relevant packages (R Core Team, 2023). The correlation analysis was performed using base R (R Core Team, 2023). The mixed model analysis was performed using ASREML-R version 4.2 (Butler et al., 2018). The trial locations were visualized on maps using the R packages ‘sf’ (Pebesma, 2018) and ‘rnaturalearth’ (Massicotte & South, 2024). The percentage change in realized genetic gain was calculated from the regression model using the *parameters_gg* function in the R package ‘agriutilities’ (https://rdrr.io/github/AparicioJohan/agriutilities/). The visualizations for descriptive statistics and genetic trends were created using the R package ‘ggplot2’ (Wickham, 2016).

## 3. Results

### 3.1 Connectivity in the historical dataset

Estimation of genetic gains using historical datasets needs appropriate connectivity over the years. The HRSWURN dataset used in this study had good connectivity based on two long-term and multiple short-term checks (Figure 4). Long-term checks (‘Marquis’ and ‘Chris’) were included from 1968 to 2023, except for 2011 and 2012. In addition to long-term checks, there were 11 short-term checks used in phases, including prominent cultivars, namely ‘Waldron’, ‘Butte’, ‘Era’, ‘Butte-86’, ‘Stoa’, ‘Keene’, ‘Verde’, ‘2375’, ‘Prosper’, ‘Linkert’, and ‘Boost’ (Figure 4). Furthermore, many of the experimental lines were tested for two or more consecutive years, ensuring stable connectivity over time for estimating genetic values for various traits. For example, we observed up to 19 shared genotypes for two consecutive years (Figure 4).

**Figure 4.**
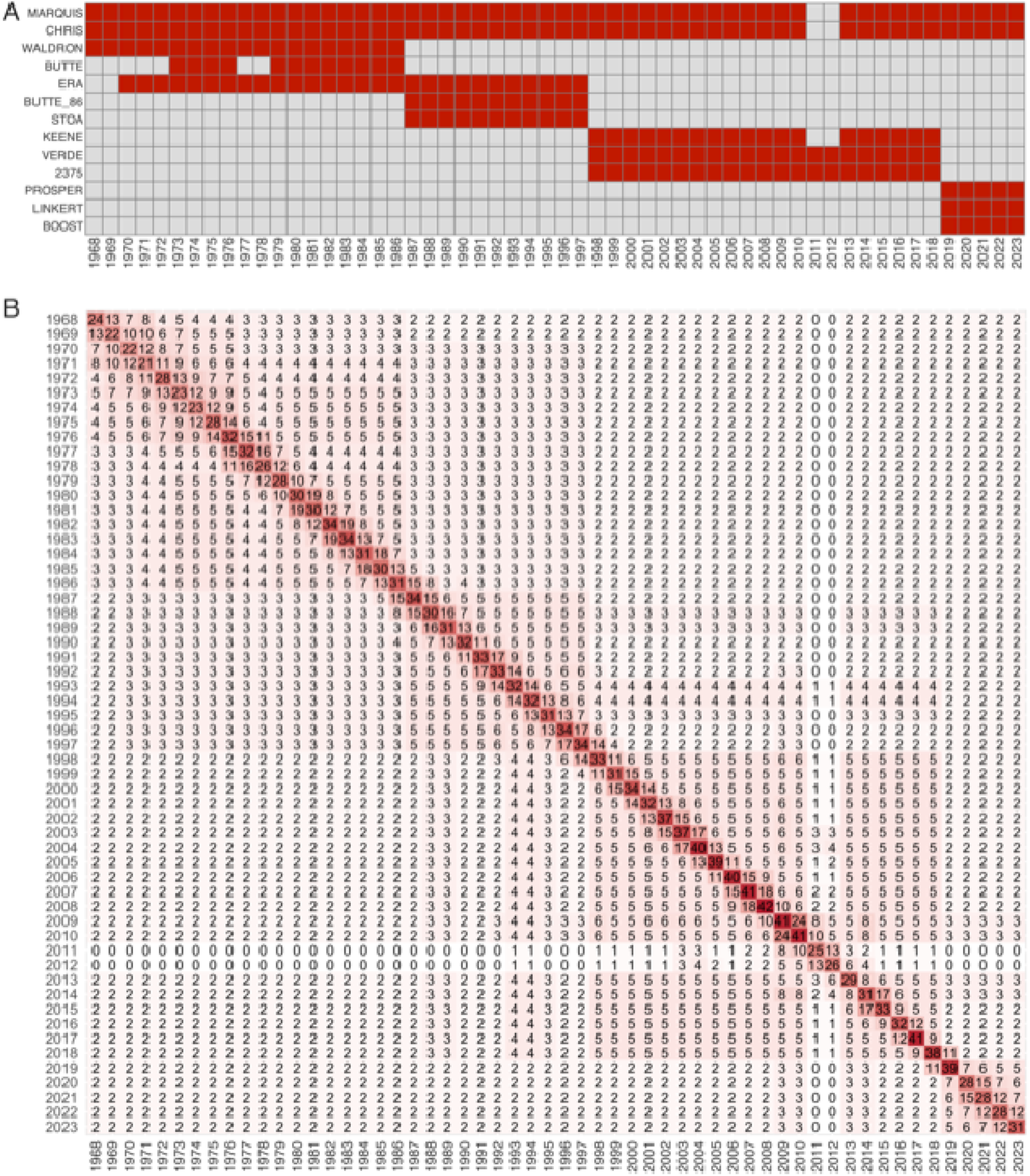
Connectivity across years from 1968 to 2023. (A) The upper panel presents the connectivity through long- and short-term checks. (B) The lower panel displays a heatmap of connectivity using the checks and genotypes tested over multiple years. The diagonal shows the number of unique genotypes tested in a given year. The off-diagonal values represent the number of common genotypes between any two given years.

### 3.2 Descriptive statistics and trait correlations

The descriptive statistics were calculated based on the genotypic BLUEs for unique genotypes obtained from the mixed-model analysis. The summary of five traits used in this study (YLD, TWT HD, HT, and PROT) is presented in Supplementary Table S3. The grain yield ranged from 2,391 kg ha^-1^ to 4,727 kg ha^-1^, with a mean value of 3,495 kg ha^-1^. The grain protein content ranged from 12.7% to 16.4%. In addition, the phenotypic differences were evident for TWT, HD, and HT (Supplementary Table S3). The trait correlations were calculated for the complete dataset based on the genotype BLUEs (Figure 5A). Grain yield (YLD) had a significant positive correlation with TWT (r = 0.51) but a significant negative correlation with PROT (r = −0.31) and HT (r = −0.33). Further, we also examined the correlations among agronomic traits in 20-year phases as the pre-1984 period (Figure 5B), 1984-2003 (Figure 5C), and 2004-2023 (Figure 5D) to study the correlation pattern over the period of adoption of semidwarf wheat. Although we observed a similar correlation pattern among most traits in different phases, the negative correlation between YLD and HT showed a continuous decline over time, from r = −0.28 in phase 1 to r = −0.11 in phase 3.

**Figure 5.**
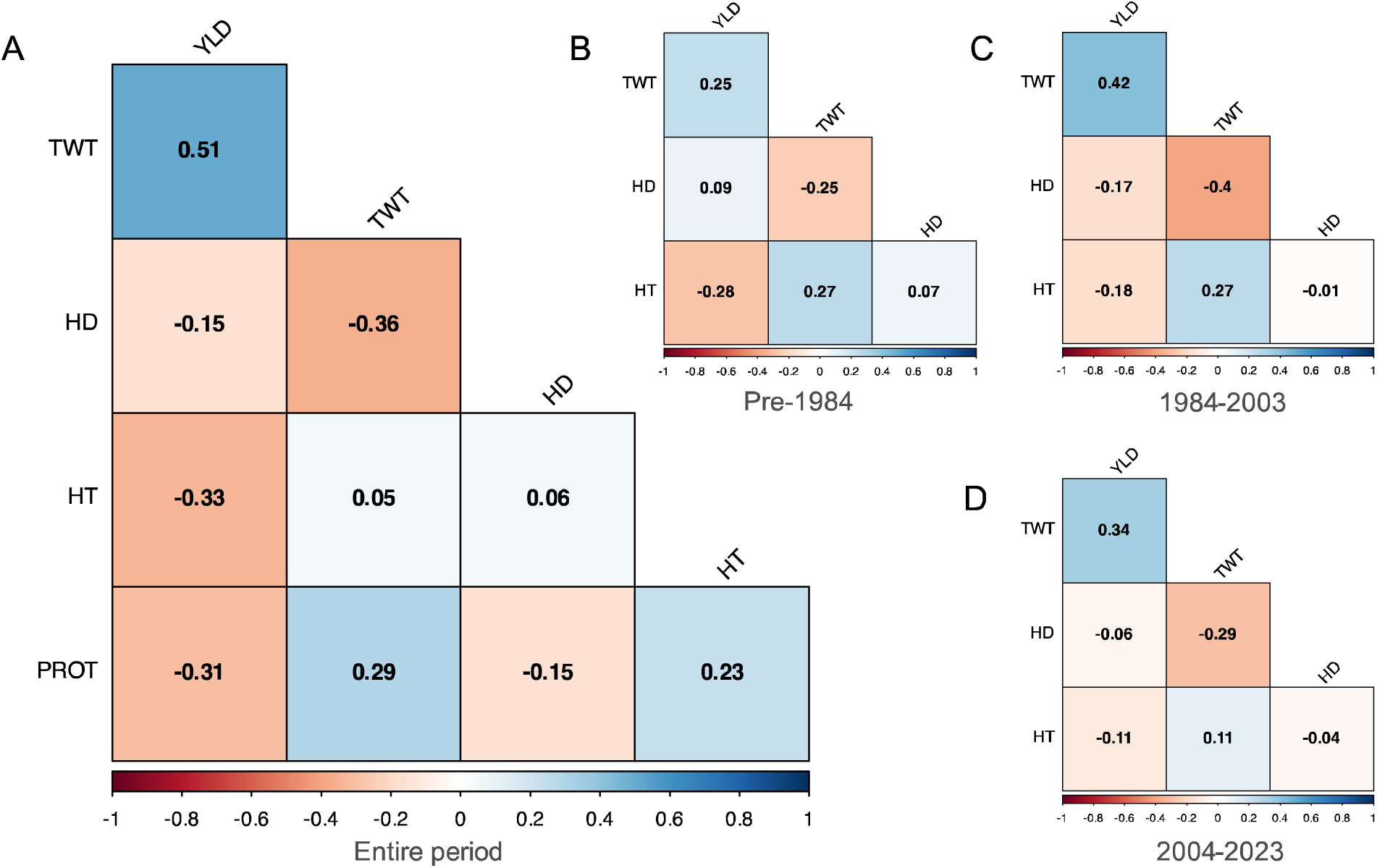
Correlation matrix presenting Pearson’s correlation among different agronomic traits in (A) the overall dataset, a subset of the complete datasets from (B) pre-1984, (C) 1984-2003, and (D) 2004-2023. The correlations with PROT in (A) are from 1995 to 2023.

### 3.3 Realized genetic gains in HRS wheat in North America

The realized gains were estimated for important agronomic traits over the past six decades in the HRS wheat from the Northern US using the HRSWURN dataset. The genetic gain was calculated separately for all tested material and released cultivars. Across the entire spring wheat region, a significant positive trend was observed for yield and test weight (Figure 6A). The absolute realized genetic gain for YLD was 15.39 kg ha^-1^ per annum, which translates to a genetic gain of 0.53% per year. The gains were higher when considering only released cultivars from the entire region (released by various programs), with an absolute realized gain of 17.68 kg ha^-1^ per year (0.61% per annum) in grain yield.

**Figure 6.**
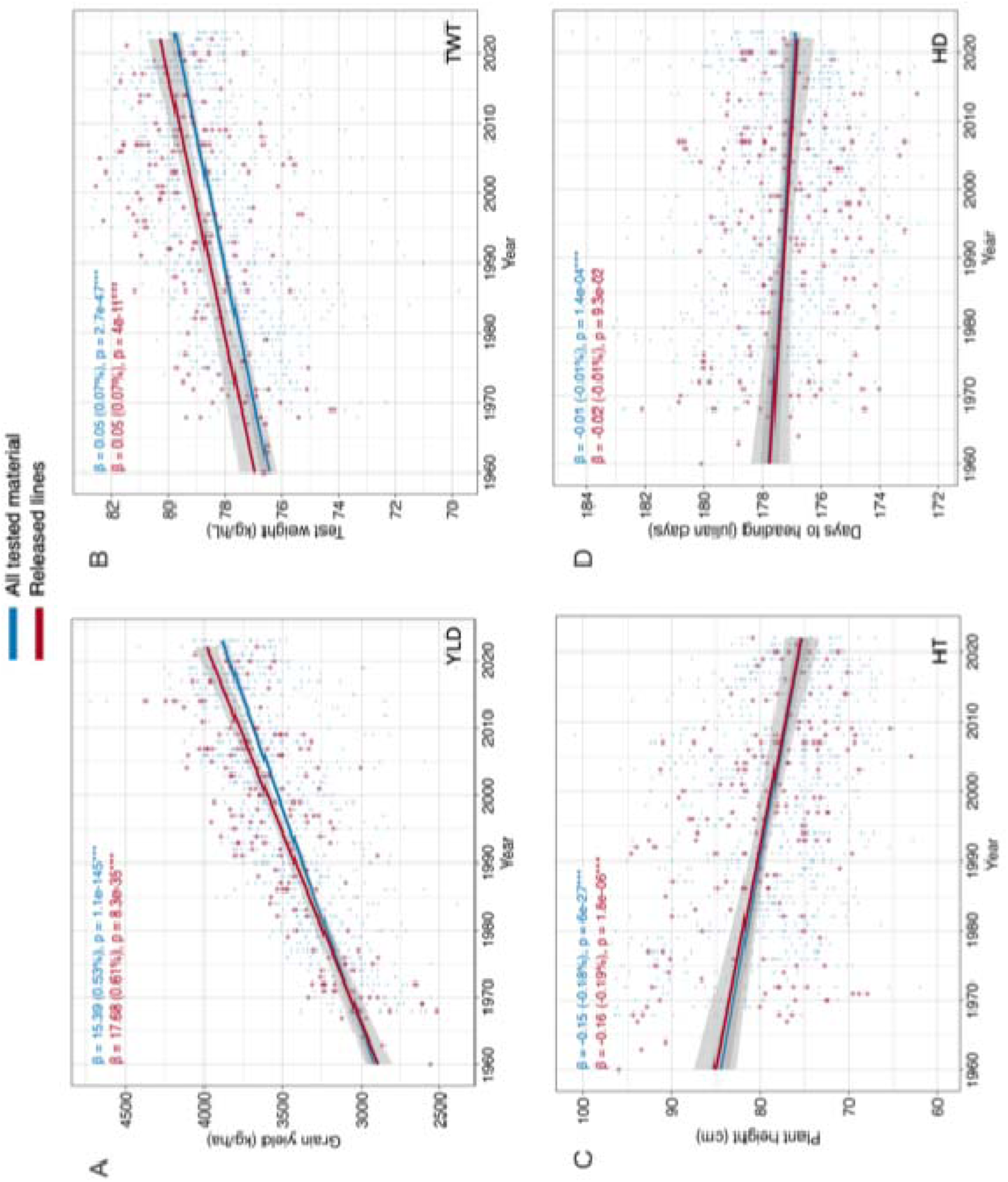
Genetic gains from breeding efforts for HRS wheat in the Northern US over six decades. The scatterplots show trends for (A) grain yield, (B) test weight, (C) plant height, and (D) days to heading. The small/blue points in the scatterplot represent all tested lines, while the large/red points represent the released cultivars. The blue line represents the trends for all the tested materials, while the red line represents the trends based only on released wheat lines (from public and private breeding programs throughout the region). The beta value (slope) from the regression equation is the absolute realized genetic gain in units per year, and its translation to percentage is provided in parentheses. Though we used the HRSWURN data from 1968, a few genotypes in the initial years were included in the HRSWURN from 1960. We used the exact year they were included in the nursery or the year they were released as a cultivar as the year of origin.

Similar positive trends were observed for TWT, with a realized genetic gain of 0.07% per annum for released cultivars (Figure 6B). For plant height, we observed a significant negative trend in all tested lines (−0.18% per year) as well as in the subset of released cultivars (−0.18% per annum) (Figure 6C). It was also evident that the variation for plant height gradually decreased over time, especially in the past two decades (Figure 6C). Further, a slightly negative trend was observed for days to heading (Figure 6D). Similar to plant height, we also observed a decline in variation for day to heading in recent decades. The data on grain protein content were only available from the mid-1990s to 2023 and were used to assess the genetic gains for this trait. Despite a strong negative correlation between YLD and PROT (*r* = −0.31), we did not observe any significant trend for PROT, with a nearly zero slope regression line for all tested materials (*p =* 0.08) and released lines (*p =* 0.09) (Figure 7).

**Figure 7.**
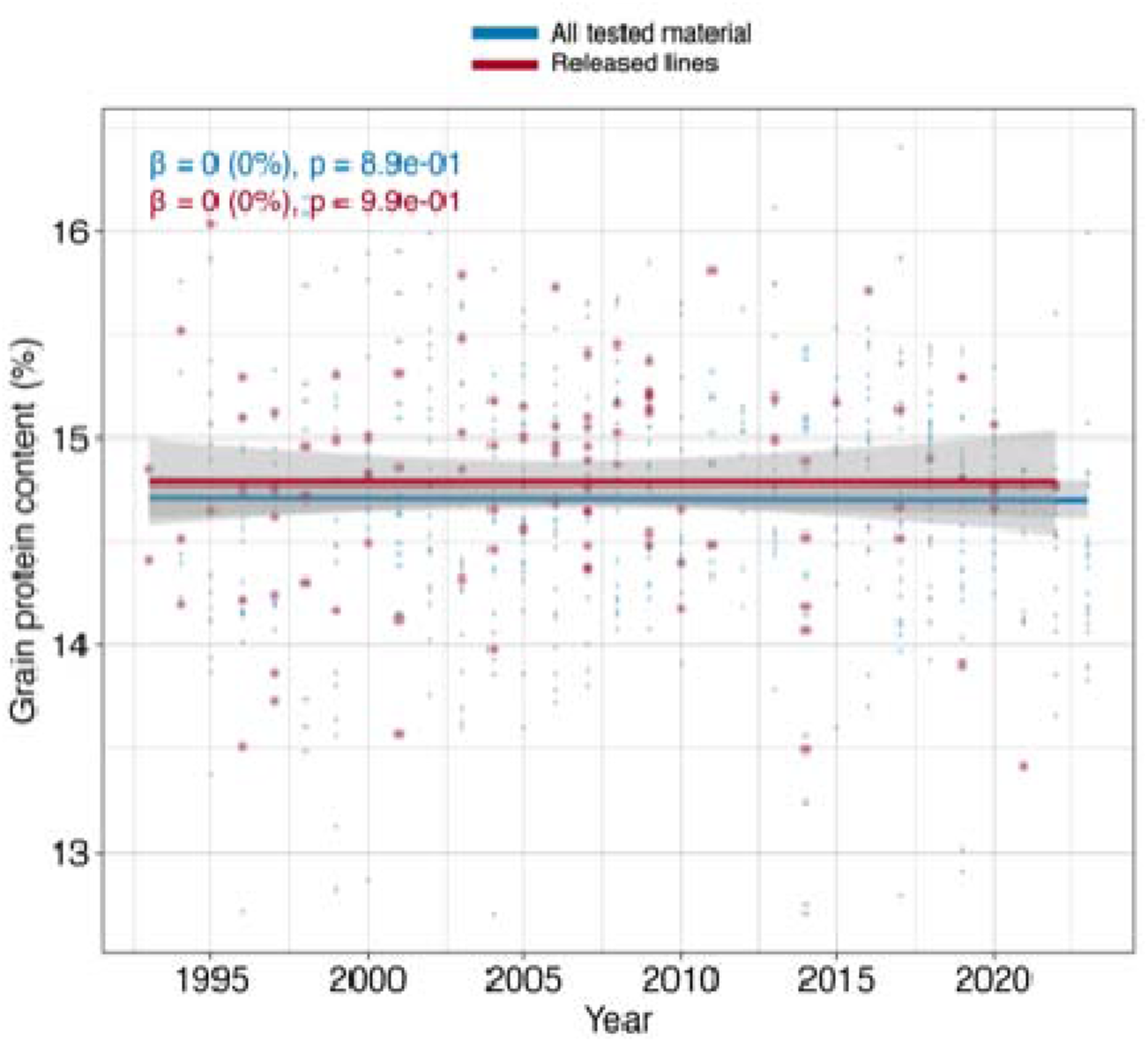
Genetic gains for grain protein content (PROT) in HRS wheat from the Northern US from the mid-1990s to 2023. The small/blue points in the scatterplot represent all tested lines, while the large/red points represent the released cultivars. The blue line in the scatterplot represents the trends for all the tested materials, while the red line represents the trends based only on released wheat lines from public and private breeding programs across the entire region. The beta value (slope) from the regression equation is the absolute realized genetic gain in units per year, and its translation to percentage is provided in parentheses.

### 3.4 Slowing rate of realized gains but no plateaus

To better understand the regional gains, we divided the entire dataset into three phases: the pre-1984 period, the second phase from 1984 to 2003, and the third phase from 2004 to 2023, with each phase representing approximately two decades of breeding efforts. In the first phase, the absolute realized gain for YLD using released cultivars was 29.68 kg ha^-1^ per year, which translates to a 1.1% increase per year (Figure 8). The second phase witnessed a decline in the genetic gains for YLD, with the absolute realized gain plummeting to 9.04 kg ha^-1^ per year (0.26% per annum). In the most recent phase, the estimated realized gains were lower than in the first phase; however, we observed an improvement over phase 2, and the absolute realized gains for YLD were 13.7 kg ha^-1^ (0.37%) per year (Figure 8).

**Figure 8.**
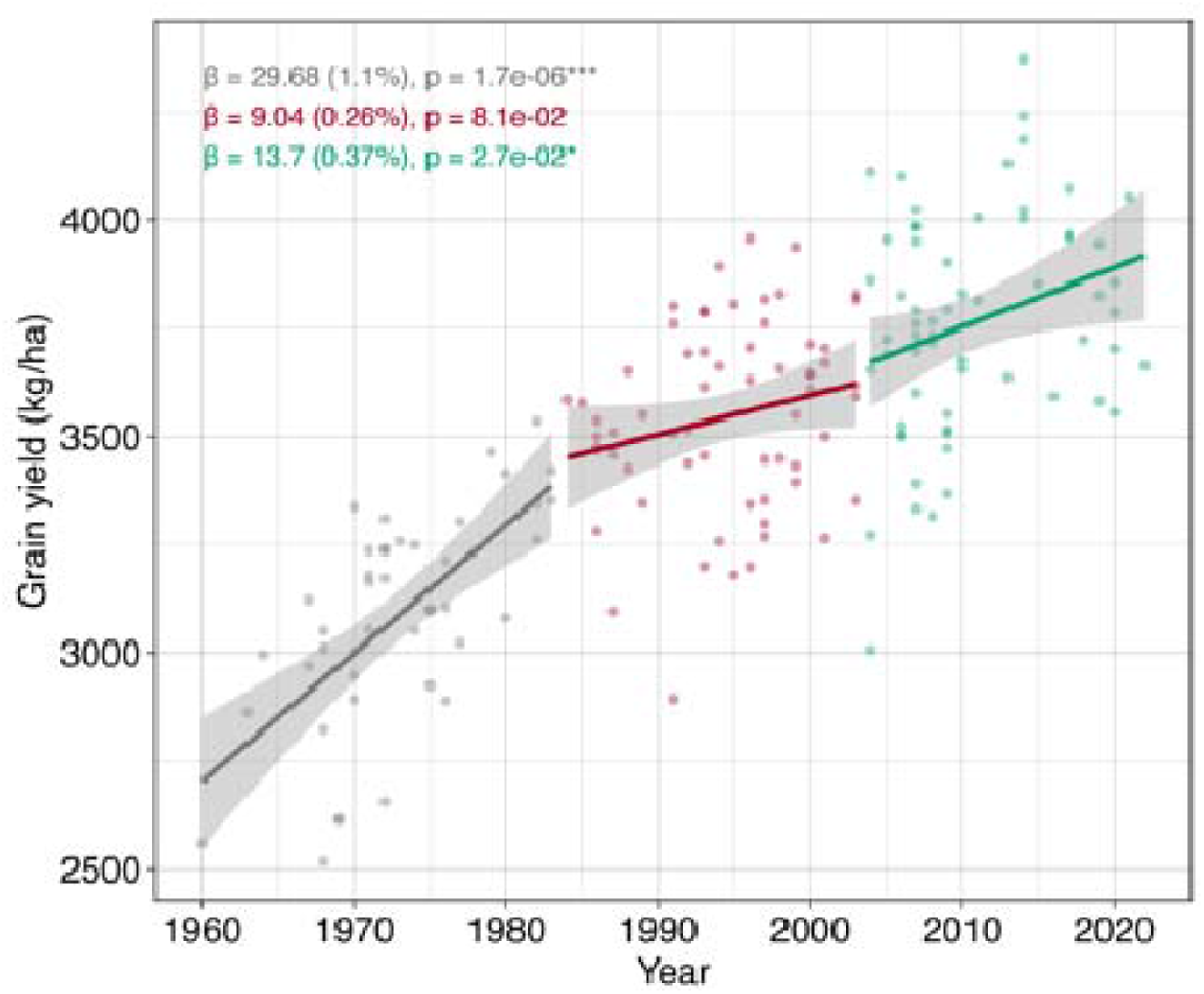
Genetic gains for grain yield in the HRS wheat from the northern region over three time periods based on the cultivars released by various organizations. The first phase spanned from 1960 to 1983, the second phase from 1984 to 2003, and the third phase from 2004 to 2023.

### 3.5 Genetic gains for grain yield from TPEs

Furthermore, we investigated the genetic gains for grain yield observed by individual public breeding programs from Minnesota (MN), South Dakota (SD), and North Dakota (ND). A new estimate of genetic gains was calculated using data from a subset of lines developed by each of these programs and evaluated in the locations within the TPEs of each program (Supplementary Table S2). A higher genetic gain for YLD was observed for MN and SD programs compared to the regional gains (Table 2). The long-term absolute realized gain in the MN breeding program was 29.20 kg ha^-1^ per year, translating to annual gains of 0.98 percent.

**Table 2.** Realized genetic gains for grain yield for the individual breeding programs based on the analysis from their respective target population of environments. The realized gains are based on the released cultivars from individual programs.

| Program | First year in analysis | Slope (Absolute gain) | SE for Slope | Intercept | R<sup>2</sup> | Pr(>F) | Genetic Gain year<sup>-1</sup> (%) |
| --- | --- | --- | --- | --- | --- | --- | --- |
| Minnesota | 1968 | 29.20 | 2.85 | -54485.75 | 0.80 | 8.46E-11 | 0.98 |
| South Dakota | 1976 | 20.46 | 4.09 | -37601.33 | 0.51 | 4.14E-05 | 0.72 |
| North Dakota | 1960 | 13.56 | 2.14 | -23564.21 | 0.54 | 3.05E-07 | 0.45 |

Similarly, the long-term absolute realized gain was kg ha^-1^ (0.72% per year) for the SD program, which is higher than the regional realized gain of 0.61% per year. Contrarily, the realized gains for the ND program were kg ha^-1^, which was slightly lower than the regional gain of 17.68 kg/ha per annum (Table 2).

## 4. Discussion

Wheat breeders worldwide face the challenge of accelerating genetic gains to address the growing global demand for food and feed. Predictive models indicate that breeding programs must achieve significantly higher rates of genetic progress to meet future wheat demands. To design effective strategies for achieving these gains, it is crucial to evaluate the progress made by breeding programs to date. In recent years, the estimation of genetic gain has been used as a key indicator for assessing the effectiveness of breeding efforts, analyzing the strengths and weaknesses of the program, and incorporating novel strategies adapted to new scenarios (Covarrubias-Pazaran, 2020; Seck et al., 2023).

Various approaches for estimating realized genetic gain in plant breeding have been proposed, utilizing different response variables in regression analyses. Numerous studies have successfully utilized historical datasets in wheat, particularly from coordinated regional nurseries, to quantify yield gains over different periods. In the US, USDA-coordinated hard winter wheat Northern Regional Performance Nursery (NRPN) and Southern Regional Performance Nursery (SRPN) have been regularly used to assess genetic gains at various intervals (Boehm et al., 2023; Graybosch & Peterson, 2010; Rife et al., 2019; Schmidt & Worrall, 1983). In this study, we utilized a historical dataset from HRSWURN, a long-term, coordinated testing platform in the Northern US, to assess regional genetic gains for hard red spring wheat. This assessment was necessary because no previous study had quantified regional gains for HRS from this region. The HRSWURN dataset has high TPE coverage for the entire region, as well as for individual breeding programs participating in the nursery. The dataset has been well-connected over the years through long-term and short-term checks (Figure 4). As recommended by Raymond et al. (2022), including at least two long-term checks can help reduce estimation bias when fitting a linear mixed model to estimate genetic gains. It is noteworthy that the susceptibility of two long-term checks (‘Marquis’ and ‘Chris’) to diseases such as wheat rusts and FHB could affect the adjustment of the year effect when calculating BLUEs. However, it’s been rare that leaf or stem rust significantly affects yields - the infections are often later in the season and don’t develop fast enough to have much effect because the wheat is near senescence by that time. Additionally, we excluded several locations from the analysis based on specific remarks in the metadata, such as those affected by the FHB or rust epidemics. Furthermore, the inclusion of short-term checks and year-to-year connectivity helps to alleviate this problem.

In this study, the realized genetic gain for grain yield was 0.61% per year for the entire North American HRS wheat region, which is comparable to long-term genetic gains in wheat improvement programs from various regions of the world. For instance, Gummadov et al. (2015) reported a realized genetic gain of 0.65% per year in wheat yield resulting from breeding efforts in Turkey from 1963 to 2004. Similarly, realized genetic gains for grain yield have been reported to be 0.54% in major wheat-growing regions of India from 1900 to 2016 (Yadav et al., 2021), 0.88% in Spain from 1930 to 2000 (Sanchez-Garcia et al., 2013), and 0.70% in Siberia from 1900 to 1997 (Morgounov et al., 2010). The realized gains from two different wheat-growing regions in Northern China were 0.48% per year from 1964 to 1995 and from 1960 to 2000 (Zhou et al., 2007).

In the US, several studies have reported realized genetic gains in wheat using era trials or historical trials, but the majority of them have been limited to the hard red winter wheat region of the Great Plains (Battenfield et al., 2013; Berzonsky & Lafever, 1993; Boehm et al., 2023; Fufa et al., 2005; Graybosch & Peterson, 2010; Rife et al., 2019). Most of these studies reported genetic gains over the mean yield of ‘Kharkof’, a historical check cultivar released in 1919. For comparison, the regional annual yield gain of 0.61% from this study translates to an annual gain of 0.72% over the historical cultivar Marquis. Earlier studies from the hard winter region reported realized genetic gains of 1% for 1919-1987 (Cox et al., 1988), 0.44% for 1919-1996 (Donmez et al., 2001), 0.82% for 1959-2008 (Graybosch & Peterson, 2010), and 0.40% for 1971-2008 (Battenfield et al., 2013). Recently, Boehm et al., (2023) used a regionally coordinated nursery, similar to HRSWURN, to quantify the yield gains for hard red winter wheat from the Central US from 1959 to 2021, very similar to the period utilized in our study. They reported a realized genetic gain of 0.67%, which is slightly lower than the 0.72% gain observed in our study. In addition to hard red winter wheat, Berzonsky & Lafever, (1993) observed an annual genetic gain of 0.55% for soft red winter wheat cultivars released from 1871 to 1987. Overall, the realized genetic gain for HRS wheat from the Northern US is comparable to other classes of wheat grown in different regions of the country.

Previous studies from North America, particularly from the Northern and Southern Great Plains, have suggested a decline in the relative rate of yield gains. Graybosch and Peterson (2010) analyzed the historical hard red winter wheat SRPN data and reported a significant linear trend from 1959 to 1984, followed by a non-significant trend in wheat yields over the time period 1984 to 2008. A similar observation was evident from a recent analysis of the historical NRPN dataset, showing a linear increase in grain yield from 1959 to 2008, followed by stagnation in yield gains over the recent decade (Boehm et al., 2023). In that study, genetic gains were assessed in three different phases representing time periods of 1959 to1984, 1984 to 2008, and 2008 to 2021, with absolute yield gains of 47.0, 61.7, and 8.9 kg ha^-1^ per year, respectively, with the authors interpreting that winter wheat adapted to the Northern Great Plains may have reached a yield plateau and could be approaching their potential ceiling. In our study, we observed a different pattern with the highest realized gains from 1960 to 1983 (1.1%), a non-significant but positive trend for 1984 to 2003, followed by a significant positive trend from 2004 to 2023 (Figure 8), which clearly suggests that the relative rate of yield gain has declined but not a yield plateau for HRS wheat from Northern America as reported previously (R. A. Fischer & Edmeades, 2010). In our results, the high genetic gains in the first phase reflect the impact of introducing semidwarf wheat and its early adoption in the HRS region by several breeding programs. The reduced genetic gain for yield reflected in our results during 1984 to 2003 can likely be attributed to the devastating Fusarium head blight (FHB) epidemics of the 1990s, which caused unprecedented damage to spring wheat production in the Northern Great Plains. The absence of resistance in adapted germplasm at the time prompted breeders to shift their focus toward identifying and integrating exotic resistance genes. For example, the quantitative trait locus *Fhb1* (Liu et al., 2006), originally introduced from the cultivar ‘Sumai 3’, was subsequently incorporated into locally adapted cultivars to enhance FHB resistance (McMullen et al., 2012). These efforts may have indirectly contributed to a decline in yield progress, as evident in the results. In the third phase, spanning 2004 to 2023, a significant trend in grain yield was observed (Figure 8), which could be attributed to the broad incorporation of FHB resistance-adapted material, early efforts in molecular breeding, as well as the availability of genomics in the last decade. A concerning pattern was observed in recent years, particularly after 2015, indicating a stagnation in grain yields. However, the 2010-2015 period included some higher-yielding lines and varieties that tended to have lower end-use quality. Post-2015, there may have been greater emphasis on lodging resistance and quality, which might have led to a slight decline in yield under these selection pressures. Additionally, extreme droughts in the region were another factor contributing to lower mean yields over the past five years. Thus, this short time window is insufficient to determine whether grain yields exhibit stagnation in the long term.

In addition to the grain yield, we observed a significant negative trend for plant height and a non-significant but negative trend for days to heading, which agrees with studies from various wheat classes (Berzonsky & Lafever, 1993; Boehm et al., 2023; Cox et al., 1988). A positive trend was observed for test weight with consistently higher gain for released cultivars, suggesting that test weight has been an important trait in selection (Figure 6B). Furthermore, our results indicate that breeders have successfully maintained the protein content and end-use quality despite a continuous increase in grain yield. However, we observed a lower range for grain protein content over the past few years (Figure 7), which could be worrisome, and breeders need to maintain enough genetic variation for protein content.

In addition to assessing regional gains for HRS wheat, we analyzed the genetic gains observed in three major public breeding programs in Minnesota, South Dakota, and North Dakota. We found differences in yield gains across the programs, as observed in previous studies (Rife et al., 2019; Sanchez-Garcia et al., 2013; Zhou et al., 2007). The realized genetic gains for grain yield were notably higher in the Minnesota (0.98% per year) and South Dakota (0.72% per year) programs compared to the regional gains (0.61% per year), corroborating studies that report higher gains when breeding is tailored to specific target environments (Delgado et al., 2024; Khalil et al., 2002). In contrast, the North Dakota breeding program exhibited lower realized gains for grain yield (0.45% per year). The higher gains in Minnesota and South Dakota can be attributed to their more homogeneous, concentrated spring wheat production areas. In Minnesota, production is primarily focused in the northwestern and some west-central counties, while in South Dakota, it is concentrated in the north-central, northeast, and northwest regions. These relatively smaller and well-defined target population environments enable the breeding programs to optimize selection for these specific areas, leading to improved genetic gains. Conversely, North Dakota’s TPE is much larger and more geographically dispersed, posing greater challenges for achieving higher genetic gains, as varieties must be developed to perform across a broader range of environments. Additionally, the higher gains in the Minnesota breeding program can also be attributed to the early adoption of semi-dwarf wheat compared to other breeding programs.

A similar study (Underdahl et al., 2008) was performed to assess the genetic gains using HRS wheat cultivars released by the North Dakota program from 1968 to 2006 based on an era trial, and the realized gain for grain yield was 1.3% per year, which is much higher than the realized gain observed in the current study. The differences in these two studies could be attributed to two main reasons. First, the era trial was conducted at only three locations, concentrated in a single region of the state, and thus did not represent the TPEs. Secondly, the era study consisted of 33 genotypes released across 40 years, and the trial was not protected by fungicides that could affect the performance of older cultivars. As suggested by Battenfield et al. (2013), this scenario can favor the newer genotypes with better disease-resistance packages, especially for Fusarium head blight in this case, and may result in higher genetic gains. This also exemplifies the differences in genetic gain estimates due to different approaches, suggesting further research on methods to minimize bias in estimates.

Furthermore, we utilized historical on-farm yield trial data from Minnesota to contextualize the genetic gains observed in our study and to assess the extent to which genetic improvements from breeding programs translate into increased farm yields. The annual yield gain over time was estimated by fitting an independent linear model to the annual yield data from 1968 to 2022, based on data obtained from the USDA National Agricultural Statistics Service (https://quickstats.nass.usda.gov/; Figure 9). The analysis revealed an annual yield increase of 34 kg/ha (∼1.7% per year), compared to 29.20 kg/ha per year (∼1% per year) of realized genetic gain in Minnesota during the same period. This higher on-farm yield increase suggests the transfer of improved genetics to growers’ fields and also highlights the synergistic effects of genetic improvement and agronomic innovations, including better cultural practices, agrochemicals, and farm mechanization, leading to much higher on-farm yield gains. These results underscore the importance of integrating genetic improvements with cutting-edge agronomic practices to maximize yield potential and ensure sustainable wheat production that meets future needs.

**Figure 9.**
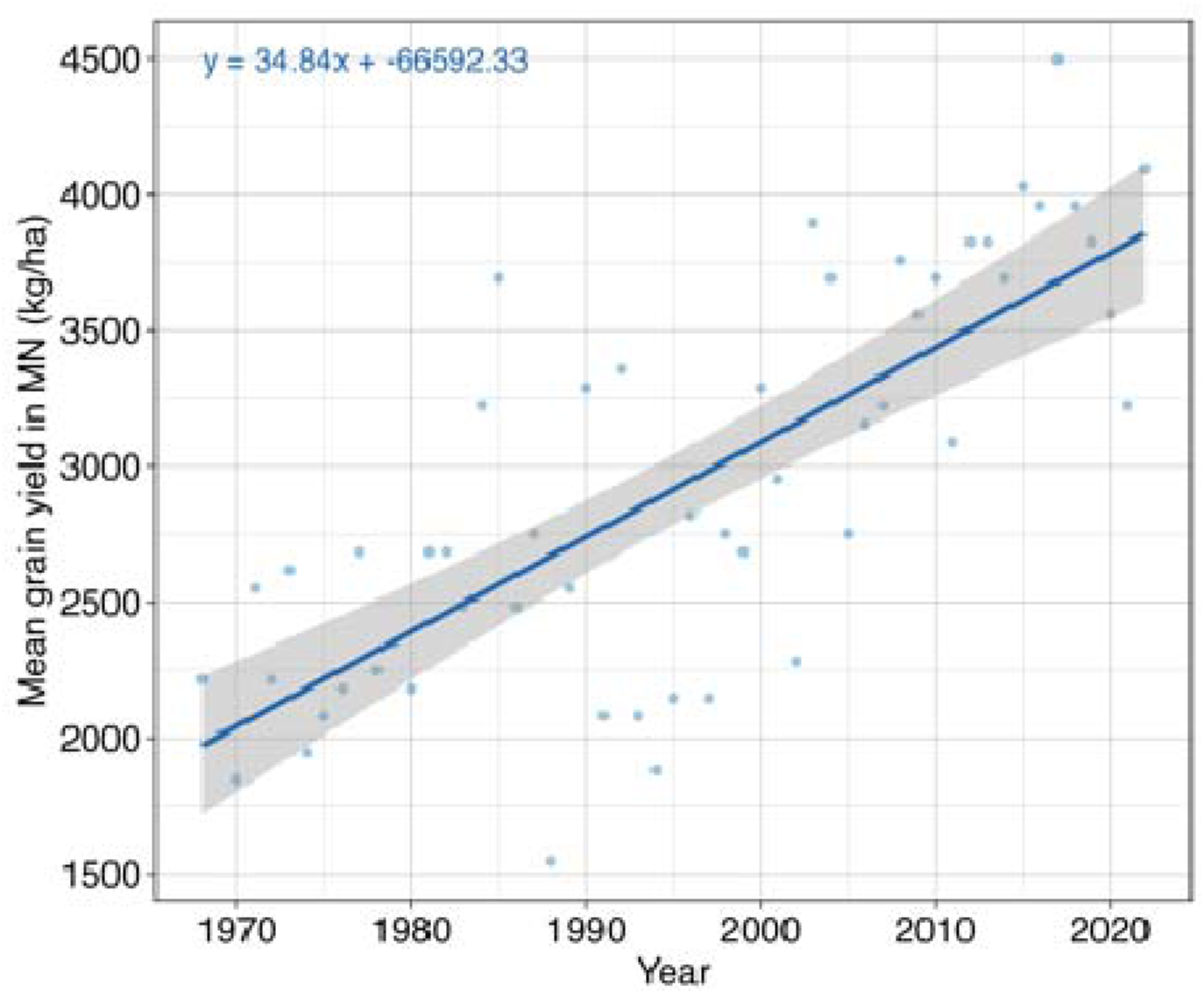
On-farm spring wheat yield trends in Minnesota from 1968 onwards. The slope of the regression equation represents the annual increase in grain yield over time.

In conclusion, these findings highlight the substantial and long-term impact of breeding efforts in the Northern Great Plains, providing important information for refining future strategies to optimize breeding programs. The study also highlights the important role of public breeding programs in regional crop improvement and justifies the need for continuous funding support for regional breeding programs. Furthermore, the results indicate that the relative rates of yield increase have declined in recent decades and are significantly lower than the yield gains necessary for sustainable wheat production. Recent models have revealed that winter wheat is more resilient, but spring wheat from North America would be more prone to environmental changes and would suffer greater yield losses with temperature increases (Zhang et al., 2022), which further exacerbates the situation. A comprehensive approach, including advancing our understanding of the genetics underlying agronomic and quality traits, modernizing breeding programs, and integrating innovative technologies like genomic selection and gene editing, could be essential to ensure sustained progress in wheat breeding. By leveraging these advancements, breeding programs can accelerate genetic gains and develop resilient, high-performing wheat varieties that are well-suited for future environments.

## Acknowledgment and Funding

The authors would like to thank the many researchers who have collaborated in the HRSWURN over the past decades. We are also grateful to Dr. Jessica Rutkoski from the University of Illinois for suggestions regarding estimating genetic gains. This project was supported by the United States Department of Agriculture-Agricultural Research Service Cooperative Research Agreement No. 58-5062-3-008 and project 5062-30100-001-000D. The U.S. Department of Agriculture (USDA) prohibits discrimination in all its programs and activities on the basis of race, color, national origin, age, disability, and where applicable, sex, marital status, familial status, parental status, religion, sexual orientation, genetic information, political beliefs, reprisal, or because all or part of an individual’s income is derived from any public assistance program. (Not all prohibited bases apply to all programs.) Persons with disabilities who require alternative means for communication of program information (Braille, large print, audiotape, etc.) should contact USDA’s TARGET Center at (202) 720-2600 (voice and TDD). To file a complaint of discrimination, write to USDA, Director, Office of Civil Rights, 1400 Independence Avenue, S.W., Washington, D.C. 20250-9410, or call (800) 795-3272 (voice) or (202) 720-6382 (TDD). USDA is an equal opportunity provider and employer.

## Data Availability Statement

The datasets generated in this study are provided within the manuscript, its supplementary files, or the repositories described in the paper. The data for HRSWURN is available at https://wheat.pw.usda.gov/GG3/germplasm. The corresponding authors should be contacted for any additional requests.

## References

Asfaw, A., Agre, P. A., Dieng, I., Adebola, P., Obidiegwu, J. E., Chamba, E., Darkwa, K., Otoo, E., Dansi, A., Dibi, K. E. B., Kouakou, A. M., & Asiedu, R. (2024). Trends in genetic gain for yam in the IITA breeding program. Crop Science, 64(4), 2261–2273. 10.1002/CSC2.21289

Ayenew, B. Z., Dejene, T., & Worede, F. (2021). Genetic Gain in Yield and Yield Attributing Traits of Rice under Upland Ecosystem of Fogera, Northwest Ethiopia. Black Sea Journal of Agriculture, 4(2), 79–87. 10.47115/BSAGRICULTURE.639422

Battenfield, S. D., Klatt, A. R., & Raun, W. R. (2013). Genetic Yield Potential Improvement of Semidwarf Winter Wheat in the Great Plains. Crop Science, 53(3), 946–955. 10.2135/CROPSCI2012.03.0158

Berzonsky, W. A., & Lafever, H. N. (1993). Progress in Ohio soft red winter wheat breeding: Grain yield and agronomic traits of cultivars released from 1871 to 1987. Crop Science, 33(6), 1382–1386.

Boehm, J. D., Masterson, S., Palmer, N., Cai, X., & Miguez, F. (2023). Genetic improvement of winter wheat (Triticum aestivum L.) grain yield in the Northern Great Plains of North America, 1959–2021. Crop Science, 63(6), 3236–3249. 10.1002/CSC2.21065

Brancourt-Hulmel, M., Doussinault, G., Lecomte, C., Bérard, P., Buanec, B. L., & Trottet, M. (2003). Genetic Improvement of Agronomic Traits of Winter Wheat Cultivars Released in France from 1946 to 1992. Crop Science, 43(1), 37–45. 10.2135/CROPSCI2003.3700

Butler, D. G., Cullis, B. R., Gilmour, A. R., Gogel, B. J., & Thompson, R. (2018). ASReml-R Reference Manual Version 4. http://www.homepages.ed.ac.uk/iwhite/asreml/uop.

Ceccarelli, S. (2015). Efficiency of Plant Breeding. Crop Science, 55(1), 87–97. 10.2135/CROPSCI2014.02.0158

Covarrubias-Pazaran, G. E. (2020). Genetic gain as a high-level key performance indicator. Genetic Gain as a High-Level Key Performance Indicator.

Cox, T. S., Shroyer, J. P., Ben-Hui, L., Sears, R. G., & Martin, T. J. (1988). Genetic Improvement in Agronomic Traits of Hard Red Winter Wheat Cultivars 1919 to 1987. Crop Science, 28(5), 756–760. 10.2135/CROPSCI1988.0011183X002800050006X

Crespo-Herrera, L. A., Crossa, J., Huerta-Espino, J., Vargas, M., Mondal, S., Velu, G., Payne, T. S., Braun, H., & Singh, R. P. (2018). Genetic Gains for Grain Yield in CIMMYT’s Semi-Arid Wheat Yield Trials Grown in Suboptimal Environments. Crop Science, 58(5), 1890–1898. 10.2135/CROPSCI2018.01.0017

Delgado, L. F., Moreta, D. E., Morante, N., Lenis, J. I., Aparicio, J. S., Londoño, L. F., Salazar, S. M., Tran, T., Ospina, M. A., & Melendez, J. L. L. (2024). Assessing realized genetic gains in biofortified cassava breeding for over a decade: Enhanced nutritional value and agronomic performance. Crop Science.

Delgado, L. F., Moreta, D. E., Morante, N., Lenis, J. I., Aparicio, J. S., Londoño, L. F., Salazar, S. M., Tran, T., Ospina, M. A., Melendez, J. L. L., Alzate, J. L. M., Vargas, H. C., Duran, L. P., Alpala, E. A. R., & Zhang, X. (2024). Assessing realized genetic gains in biofortified cassava breeding for over a decade: Enhanced nutritional value and agronomic performance. Crop Science, 64(6), 3242–3258. 10.1002/csc2.21369

Donmez, E., Sears, R. G., Shroyer, J. P., & Paulsen, G. M. (2001). Genetic gain in yield attributes of winter wheat in the Great Plains. Crop Science, 41(5), 1412–1419. 10.2135/CROPSCI2001.4151412X

Duvick, D. N. (2005). The Contribution of Breeding to Yield Advances in maize (Zea mays L.). Advances in Agronomy, 86, 83–145. 10.1016/S0065-2113(05)86002-X

Eberhart, S. A. (1964). Least Squares Method for Comparing Progress Among Recurrent Selection Methods1. Crop Science, 4(2), 230–231. 10.2135/CROPSCI1964.0011183X000400020036X

Fischer, R. A., & Edmeades, G. O. (2010). Breeding and cereal yield progress. Crop Science, 50. 10.2135/cropsci2009.10.0564

Fischer, R., Byerlee, D., & Edmeades, G. (2014). Crop yields and global food security. In Academia.edu. ACIAR: Canberra, ACT.

Fufa, H., Baenziger, P. S., Beecher, B. S., Graybosch, R. A., Eskridge, K. M., & Nelson, L. A. (2005). Genetic improvement trends in agronomic performances and end-use quality characteristics among hard red winter wheat cultivars in Nebraska. Euphytica, 144(1–2), 187–198. 10.1007/S10681-005-5811-X

Gerard, G. S., Crespo-Herrera, L. A., Crossa, J., Mondal, S., Velu, G., Juliana, P., Huerta-Espino, J., Vargas, M., Rhandawa, M. S., Bhavani, S., Braun, H., & Singh, R. P. (2020). Grain yield genetic gains and changes in physiological related traits for CIMMYT’s High Rainfall Wheat Screening Nursery tested across international environments. Field Crops Research, 249, 107742. 10.1016/J.FCR.2020.107742

Graybosch, R. A., & James Peterson, C. (2012). Specific adaptation and genetic progress for grain yield in Great Plains hard winter wheats from 1987 to 2010. Crop Science, 52(2), 631–643. 10.2135/CROPSCI2011.08.0412

Graybosch, R. A., & Peterson, C. J. (2010). Genetic improvement in winter wheat yields in the Great Plains of North America, 1959-2008. Crop Science, 50(5), 1882–1890. 10.2135/CROPSCI2009.11.0685

Gummadov, N., Keser, M., Akin, B., Cakmak, M., Mert, Z., Taner, S., Ozturk, I., Topal, A., Yazar, S., & Morgounov, A. (2015). Genetic gains in wheat in Turkey: Winter wheat for irrigated conditions. The Crop Journal, 3(6), 507–516. 10.1016/J.CJ.2015.07.007

Khalil, I. H., Carver, B. F., Krenzer, E. G., MacKown, C. T., & Horn, G. W. (2002). Genetic trends in winter wheat yield and test weight under dual-purpose and grain-only management systems. Crop Science, 42(3), 710–715.

Khanna, A., Anumalla, M., Ramos, J., Cruz, M. T. S., Catolos, M., Sajise, A. G., Gregorio, G., Dixit, S., Ali, J., Islam, M. R., Singh, V. K., Rahman, M. A., Khatun, H., Pisano, D. J., Bhosale, S., & Hussain, W. (2024). Genetic gains in IRRI’s rice salinity breeding and elite panel development as a future breeding resource. Theoretical and Applied Genetics, 137(2), 1–14. 10.1007/S00122-024-04545-9/FIGURES/1

Kumar, A., Raman, A., Yadav, S., Verulkar, S. B., Mandal, N. P., Singh, O. N., Swain, P., Ram, T., Badri, J., Dwivedi, J. L., Das, S. P., Singh, S. K., Singh, S. P., Kumar, S., Jain, A., Chandrababu, R., Robin, S., Shashidhar, H. E., Hittalmani, S., … Piepho, H. P. (2021). Genetic gain for rice yield in rainfed environments in India. Field Crops Research, 260, 107977. 10.1016/J.FCR.2020.107977

Liu, S., Zhang, X., Pumphrey, M. O., Stack, R. W., Gill, B. S., & Anderson, J. A. (2006). Complex microcolinearity among wheat, rice, and barley revealed by fine mapping of the genomic region harboring a major QTL for resistance to Fusarium head blight in wheat. Functional & Integrative Genomics, 6(2), 83–89. 10.1007/s10142-005-0007-y

Mackay, I., Horwell, A., Garner, J., White, J., McKee, J., & Philpott, H. (2011). Reanalyses of the historical series of UK variety trials to quantify the contributions of genetic and environmental factors to trends and variability in yield over time. Theoretical and Applied Genetics, 122(1), 225–238. 10.1007/S00122-010-1438-Y/FIGURES/10

Massicotte, P., & South, A. (2024). rnaturalearth: World Map Data from Natural Earth. [Computer software]. https://github.com/ropensci/rnaturalea

McMullen, M., Bergstrom, G., De Wolf, E., Dill-Macky, R., Hershman, D., Shaner, G., & Van Sanford, D. (2012). A unified effort to fight an enemy of wheat and barley: Fusarium head blight. Plant Disease, 96(12), 1712–1728. 10.1094/PDIS-03-12-0291-FE

Menkir, A., Dieng, I., Meseka, S., Bossey, B., Mengesha, W., Muhyideen, O., Riberio, P. F., Coulibaly, M., Yacoubou, A. M., Bankole, F. A., Adu, G. B., & Ojo, T. (2022). Estimating genetic gains for tolerance to stress combinations in tropical maize hybrids. Frontiers in Genetics, 13. 10.3389/FGENE.2022.1023318

Morgounov, A., Zykin, V., Belan, I., Roseeva, L., Zelenskiy, Y., Gomez-Becerra, H. F., Budak, H., & Bekes, F. (2010). Genetic gains for grain yield in high latitude spring wheat grown in Western Siberia in 1900-2008. Field Crops Research, 117(1), 101–112. 10.1016/J.FCR.2010.02.001

Paulsen, G. M., & Shroyer, J. P. (2008). The early history of wheat improvement in the Great Plains. Agronomy Journal, 100, S-70.

Pebesma, E. (2018). Simple features for R: Standardized support for spatial vector data. R Journal, 10(1), 439–446. 10.32614/RJ-2018-009

Piepho, H. P., Laidig, F., Drobek, T., & Me Yer, U. (2014). Dissecting genetic and non!Zlgenetic sources of long!Zlterm yield trend in german official variety trials. Theoretical and Applied Genetics, 127(5), 1009–1018. 10.1007/S00122-014-2275-1

Prasanna, B. M., Burgueño, J., Beyene, Y., Makumbi, D., Asea, G., Woyengo, V., Tarekegne, A., Magorokosho, C., Wegary, D., Ndhlela, T., Zaman-Allah, M., Matova, P. M., Mwansa, K., Mashingaidze, K., Fato, P., Teklewold, A., Vivek, B. S., Zaidi, P. H., Vinayan, M. T., … Cairns, J. E. (2022). Genetic trends in CIMMYT’s tropical maize breeding pipelines. Scientific Reports, 12(1). 10.1038/S41598-022-24536-4

R Core Team. (2023). R: A language and environment for statistical computing. [Computer software]. R Foundation for Statistical Computing. http://www.r-project.org/

Raymond, J., Mackay, I., Penfield, S., Lovett, A., Philpott, H., & Dorling, S. (2023). Continuing genetic improvement and biases in genetic gain estimates revealed in historical UK variety trials data. Field Crops Research, 302, 109086. 10.1016/J.FCR.2023.109086

Rife, T. W., Graybosch, R. A., & Poland, J. A. (2019). A Field-Based Analysis of Genetic Improvement for Grain Yield in Winter Wheat Cultivars Developed in the US Central Plains from 1992 to 2014. Crop Science, 59(3), 905–910. 10.2135/CROPSCI2018.01.0073

Rutkoski, J. E. (2019a). A practical guide to genetic gain. Advances in Agronomy, 157, 217– 249. 10.1016/BS.AGRON.2019.05.001

Rutkoski, J. E. (2019b). Estimation of Realized Rates of Genetic Gain and Indicators for Breeding Program Assessment. Crop Science, 59(3), 981–993. 10.2135/CROPSCI2018.09.0537

Sanchez-Garcia, M., Royo, C., Aparicio, N., Martín-Sánchez, J. A., & Álvaro, F. (2013). Genetic improvement of bread wheat yield and associated traits in Spain during the 20th century. The Journal of Agricultural Science, 151(1), 105–118. 10.1017/S0021859612000330

Schmidt, J. W., & Worrall, W. D. (1983). Trends in yield improvement through genetic gains. Proceedings of the Sixth International Wheat Genetics Symposium/Edited by Sadao Sakamoto.

Seck, F., Covarrubias-Pazaran, G., Gueye, T., & Bartholomé, J. (2023). Realized Genetic Gain in Rice: Achievements from Breeding Programs. Rice, 16(1). 10.1186/S12284-023-00677-6

Singh, R. P., Huerta-Espino, J., Sharma, R., Joshi, A. K., & Trethowan, R. (2007). High yielding spring bread wheat germplasm for global irrigated and rainfed production systems. Euphytica, 157(3), 351–363. 10.1007/S10681-006-9346-6/TABLES/7

Underdahl, J. L., Mergoum, M., Ransom, J. K., & Schatz, B. G. (2008). Agronomic traits improvement and associations in hard red spring wheat cultivars released in North Dakota from 1968 to 2006. Crop Science, 48(1), 158–166.

USDA ERS. (2023). USDA Economic Reseasrch Service: Wheat Data. https://www.ers.usda.gov/data-products/wheat-data/

Van Dijk, M., Morley, T., Rau, M. L., & Saghai, Y. (2021). A meta-analysis of projected global food demand and population at risk of hunger for the period 2010–2050. Nature Food, 2(7), 494–501.

Wickham, H. (2016). ggplot2: Elegant Graphics for Data Analysis. Springer-Verlag New York. https://cran.r-project.org/web/packages/ggplot2/citation.html

Xu, Y., Li, P., Zou, C., Lu, Y., Xie, C., Zhang, X., Prasanna, B. M., & Olsen, M. S. (2017). Enhancing genetic gain in the era of molecular breeding. Journal of Experimental Botany, 68(11), 2641–2666. 10.1093/JXB/ERX135

Yadav, R., Gupta, S., Gaikwad, K. B., Bainsla, N. K., Kumar, M., Babu, P., Ansari, R., Dhar, N., Dharmateja, P., & Prasad, R. (2021). Genetic Gain in Yield and Associated Changes in Agronomic Traits in Wheat Cultivars Developed Between 1900 and 2016 for Irrigated Ecosystems of Northwestern Plain Zone of India. Frontiers in Plant Science, 12. 10.3389/FPLS.2021.719394/FULL

Zhang, T., He, Y., DePauw, R., Jin, Z., Garvin, D., Yue, X., Anderson, W., Li, T., Dong, X., & Zhang, T. (2022). Climate change may outpace current wheat breeding yield improvements in North America. Nature Communications, 13(1), 5591.

Zhou, Y., He, Z. H., Sui, X. X., Xia, X. C., Zhang, X. K., & Zhang, G. S. (2007). Genetic improvement of grain yield and associated traits in the Northern China Winter Wheat Region from 1960 to 2000. Crop Science, 47(1), 245–253. 10.2135/CROPSCI2006.03.0175

