## Supplementary Data for "Genetic Gains from Sixty Years of Spring Wheat Breeding in the Northern Plains of the US"

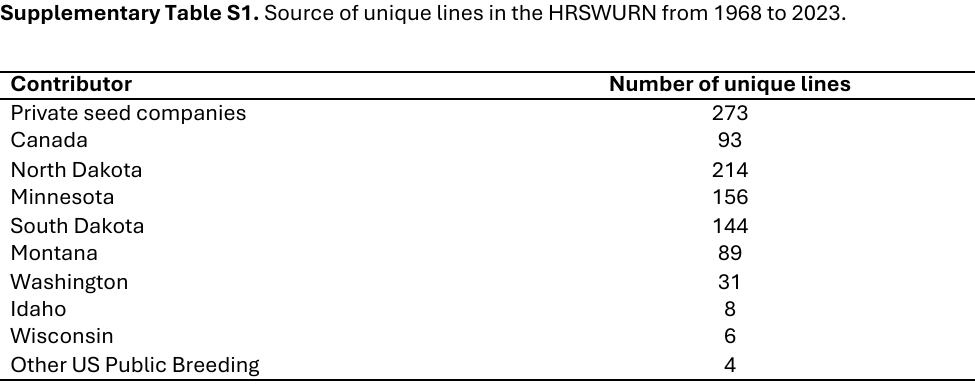

**Supplementary Table S2.** Description of the various locations across different states and provinces used for the HRSWURN trials. The TPE columns indicate if the respective location represents the target breeding environment for the public breeding program of Minnesota (TPE_MN), North Dakota (TPE_ND), or South Dakota (TPE_SD).

| **Location** | **State** | **Abbreviation** | **Latitude** | **Longitude** | **TPE_MN** | **TPE_ND** | **TPE_SD** |
| --- | --- | --- | --- | --- | --- | --- | --- |
| Aberdeen | Idaho | ABR | 42.9656285 | -112.79855 | - | - | - |
| Bonners Ferry | Idaho | BFR | 48.69211 | -116.31763 | - | - | - |
| Tetonia | Idaho | TET | 43.817389 | -111.15954 | - | - | - |
| Glenlea | Manitoba CANADA | GLL | 49.6503038 | -97.112911 | - | - | - |
| Winnipeg | Manitoba CANADA | WNP | 49.811226 | -97.12542 | - | - | - |
| Crookston | Minnesota | CRK | 47.806576 | -96.608666 | YES | YES | - |
| Morris | Minnesota | MRS | 45.6140369 | -95.734614 | YES | - | - |
| Sabin | Minnesota | SBN | 46.780034 | -96.648543 | YES | YES | - |
| St. Paul | Minnesota | STP | 44.990279 | -93.179955 | YES | - | - |
| Waseca | Minnesota | WSC | 44.077 | -93.5084 | YES | - | - |
| Bozeman | Montana | BZM | 45.668552 | -111.05245 | - | - | - |
| Conrad | Montana | CON | 48.172807 | -111.94713 | - | - | - |
| Havre | Montana | HVR | 48.540239 | -109.69091 | - | - | - |
| Moccasin | Montana | MSN | 47.0528817 | -109.90983 | - | - | - |
| Sidney | Montana | SID | 47.72727 | -104.14735 | - | - | - |
| Lind | Nebraska | LND | 46.96978 | -118.61513 | - | - | - |
| Mead | Nebraska | MED | 41.22861 | -96.48889 | - | - | - |
| Sidney | Nebraska | SDN | 41.142692 | -102.96595 | - | - | - |
| Brandon | North Dakota | BRD | 49.8679651 | -99.974062 | - | YES | - |
| Carrington | North Dakota | CRT | 47.5090572 | -99.117707 | - | YES | - |
| Casselton | North Dakota | CST | 46.888056 | -97.218285 | YES | YES | - |
| Dickinson | North Dakota | DSN | 46.874967 | -102.77587 | - | YES | - |
| Fargo | North Dakota | FRG | 46.8901035 | -96.803616 | YES | YES | - |
| Forman | North Dakota | FRM | 46.086415 | -97.636315 | - | YES | - |
| Hettinger | North Dakota | HTG | 46.010248 | -102.64536 | - | YES | - |
| Langdon | North Dakota | LGD | 48.7603192 | -98.367252 | - | YES | - |
| Minot | North Dakota | MNT | 48.2333496 | -101.29201 | - | YES | - |
| Prosper | North Dakota | PRP | 47.0013717 | -97.112242 | YES | YES | - |
| Thompson | North Dakota | TPS | 47.774861 | -97.104611 | YES | YES | - |
| Williston | North Dakota | WLT | 48.146847 | -103.62025 | - | YES | - |
| Saskatoon | Saskatchewan CANADA | SKT | 52.146973 | -106.64703 | - | - | - |
| Swift Current | Saskatchewan CANADA | SWC | 50.282358 | -107.85373 | - | - | - |
| Brookings | South Dakota | BRK | 44.320881 | -96.77264 | - | - | YES |
| Groton | South Dakota | GRT | 45.445 | -98.101496 | - | - | YES |
| Highmore | South Dakota | HMR | 44.52139 | -99.43944 | - | - | YES |
| Redfield | South Dakota | RED | 44.875145 | -98.51785 | - | - | YES |
| Selby | South Dakota | SLB | 45.502869 | -100.03346 | - | - | YES |
| Watertown | South Dakota | WAT | 44.8994 | -97.1151 | - | - | YES |
| Pullman | Washington | PLM | 46.723093 | -117.13975 | - | - | - |
| Madison | Wisconsin | MAD | 43.0761755 | -89.412508 | - | - | - |
| Powell | Wyoming | PWL | 44.776217 | -108.75889 | - | - | - |
| Sheridan | Wyoming | SHR | 44.79672 | -106.95897 | - | - | - |
| Torrington | Wyoming | TOR | 42.06246 | -104.18439 | - | - | - |

**Supplementary Table S3.** Descriptive statistics for the five agronomic traits based on the best linear unbiased estimator (BLUEs) for individual genotypes obtained from the linear mixed model.

| **Trait** | **Units** | **Mean** | **Std Dev** | **Min** | **Max** | **Skewness** | **H^2^** |
| --- | --- | --- | --- | --- | --- | --- | --- |
| YLD | kg/ha | 3494.84 | 347.35 | 2391.26 | 4727.19 | -0.31 | 0.41 |
| TWT | kg/hL | 78.46 | 1.92 | 69.75 | 82.74 | -0.61 | 0.78 |
| HD | Julian days | 177.23 | 1.88 | 171.98 | 184.53 | 0.27 | 0.56 |
| HT | cm | 78.93 | 6.97 | 59.51 | 100.97 | 0.47 | 0.90 |
| PROT | percent | 14.71 | 0.57 | 12.70 | 16.40 | -0.42 | 0.58 |

Supplementary Figure S1. Locations based on the cleaned dataset where test weight (TWT), grain protein content (PROT), plant height (PH), and days to heading (HD) were collected from 1968 to 2023. For a particular year and location, white indicates that data for YLD was not reported or the given location was not part of the nursery in the respective year. The scale provided in the legend elucidates the number of genotypes tested

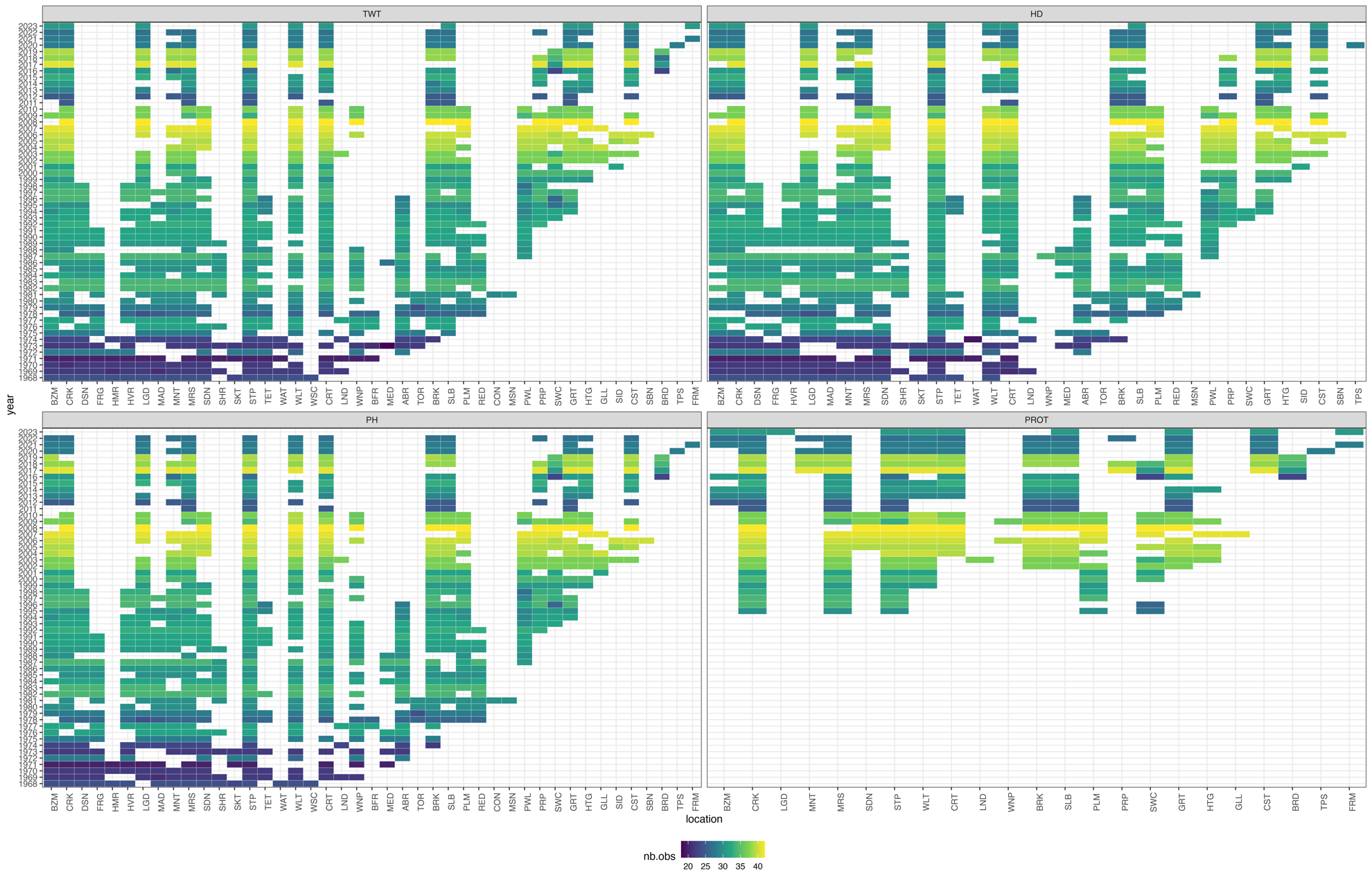
